# A UHPLC–MS/MS method for determination of N-acylethanolamines in cerebrospinal fluid

**DOI:** 10.1101/2023.11.20.567867

**Authors:** Visolakhon Ismoilova, Asger Wretlind, Kourosh Hooshmand, Anja Hviid Simonsen, Steen Gregers Hasselbalch, Cristina Legido Quigley

**Affiliations:** Institute of Cancer and Pharmaceutical Science, King’s College London, Stamford Street, SE1 9NQ, London, United Kingdom; System Medicine Group, Steno Diabetes Centre Copenhagen, Borgmester lb Juuls Vej 83, 2730 Herlev, Denmark; Danish Dementia Research Centre, Copenhagen University Hospital, Rigshospitalet, Copenhagen, Denmark

**Keywords:** N-acylethanolamines (NAEs), Primary fatty amides (PFAMs), Cerebrospinal fluid, Mass spectrometry, Lipids

## Abstract

N-acylethanolamines (NAEs) and primary fatty amides (PFAMs) are of a great interest due to the range of physiological effects they exhibit, potentially serving as neuromodulators. However, they are present at nano and picomolar concentrations in human cerebrospinal fluid (CSF) samples, posing challenges for detection and measurement using conventional Ultra-high performance liquid chromatography systems coupled to tandem mass spectrometry (UHPLC-MS).

UHPLC-MS was used in dynamic multiple reaction monitoring (dMRM) mode. Seven deuterated NAEs internal standards were used to develop the method. Six solvent combinations were tested for extraction efficiency, accuracy, precision, matrix effect, linearity, limits of detection. Lastly the method was applied to CSF from healthy individuals (n=33) to estimate their natural range of concentrations.

Extraction with acetonitrile/acetone showed the highest efficiency and recovery. The presented method was able to measure the following 17 NAEs and PFAMs in human CSF: linoleoyl ethanolamide, heptadecanoyl ethanolamide, stearoyl ethanolamide, palmitoyl ethanolamide, dihomolinolenoyl ethanolamide, eicosatrienoic acid ethanolamide, behenamide, octadecanamide, lauramide, tetradecanamide, erucamide, linoleamide, palmitamide, myristic monoethanolamide, pentadecanoyl ethanolamide, oleamide and palmitoleoyl ethanolamide. In healthy individuals the concentrations ranged three-fold from pg/mL to mg/mL. Further studies could apply this method to clinical CSF samples.

## Introduction

N-acylethanolamines (NAEs), sometimes referred to as fatty acid ethanolamides, are a narrow group of signaling lipid molecules known for their anti-inflammatory and neuroprotective roles in the brain (1– 3). Due to their role in neuro-homeostasis NAEs make for potential biomarkers for several neurological diseases, that could inform early detection, disease severity and disease progression. NAEs levels have been found to be altered in depression (4,5), brain injury (6) and Parkinson’s disease (7). Some NAEs play a role in endocannabinoid system where they can agonize the cannabinoid receptors CB1 and CB2. Anandamide (AEA) role as an endocannabinoid is well described (8–10). Docosahexaenoyl ethanolamide (DHEA) and eicosapentaenoyl ethanolamide (EPEA) have also been shown to bind the cannabinoid receptors (11).

Primary fatty amides (PFAMs) is another group of signaling molecules closely related to NAEs. The difference being a primary amide group instead of a secondary amide with an ethanol group. PFAMs have been associated to neurological dysfunction. One of the more researched PFAMs is oleamide known for its endocannabinoid and sleep-inducing properties, recently linked to Alzheimer’s (12). In this work cerebrospinal fluid (CSF) was chosen as the sample matrix best suited for measuring NAEs due to close proximation to the brain. A previous method for measuring NAEs in CSF by Kantea et al. was published using nano-LC and achieving high sensitivity for these compounds (13).

Analysis of NAEs in CSF poses some challenges. Firstly, NAEs are endogenous compounds, that exist as low-abundance metabolites in the CSF. Studies have reported presence of NAEs and their structural analogues in human CSF, with concentration ranging in the picomolar to even femtomolar concentration range (14). Adding to the complexity is the limited volume of CSF that is often collected, as the average brain CSF volume is approximately 150 mL (15). The combination of restricted sample volume and low abundance of some of these compounds poses significant challenges when it comes to identifying and quantifying the NAEs within the CSF.

NAEs present in biofluids are typically analyzed using liquid chromatography coupled to tandem mass spectrometry (LC-MS/MS) following solid-phase extraction or protein precipitation (15,16). While some study reports have used gas chromatography coupled to mass spectrometry (GC-MS) (17) to quantify the NAEs, LC is preferred due to the time-consuming derivatization step in GC-MS (15). LC-MS/MS is a highly selective and sensitive analytical approach which can be further enhanced by employing the dynamic multiple reaction monitoring (dMRM), developed to quantify the target metabolites with maximized coverage and sensitivity (18). Hence, we aimed to optimize a sensitive and robust UHPLC-MS/MS method to identify NAEs and their structural analogues PFAMs in human CSF samples. To achieve this goal, the following steps were undertaken: (i) Compare the extraction efficiency of deuterated standards in human CSF with various monophasic solvents. (ii) Assessment of the results based on the parameters including, accuracy, precision, matrix effect, linearity, limit of detection (LOD) and limit of quantification (LOQ). (iii) Measuring NAEs and PFAMs in CSF from 33 healthy individuals to know the concentration variation in healthy individuals.

## Methods

### CSF sample description

The CSF samples were collected by the Danish Dementia Research Centre, Copenhagen University Hospital, Rigshospitalet, Copenhagen, Denmark. For method development a CSF pool was prepared from 10 anonymized individuals. Human CSF samples from a cohort of 33 individuals, females (n=19) and males (n=14) were collected for identification and quantification of endogenous NAEs and PFAMs. The participants had a review of the cognitive tests, imaging and CSF analysis and were classified as cognitively healthy. The CSF samples were stored at – 80 °C until processing.

### Chemicals and reagents

Acetonitrile (ACN), formic acid (FA), isopropanol (IPA), methanol (MeOH) and ultra-pure water were purchased from Fisher Scientific (Pittsburgh, Pennsylvania, US.). Ethanol (EtOH) and acetone (Ace) were acquired from Sigma-Aldrich (Burlington, Massachusetts, US.). All solvents and reagents were of LC-MS grade and ≥98% purity. FA was used as an ionising agent, ACN, IPA and ultra-pure water were utilised for the preparation of the chromatographic mobile phases. The internal standards (ISTD) used (Table 1) were acquired from BioNordika (Herlev, Denmark).

**Table 1.** NAEs used as internal standards.

| Full Name | Abbreviation | C | C=C | HMDB |
| --- | --- | --- | --- | --- |
| Anandamide-d4 | AEA-d<sub>4</sub> | 20 | 4 | HMDB0004080 |
| Docosahexaenoyl ethanolamide-d4 | DHEA-d<sub>4</sub> | 22 | 6 | HMDB0013658 |
| Eicosapentaenoyl ethanolamide-d4 | EPEA-d<sub>4</sub> | 20 | 5 | HMDB0013649 |
| Linoleoyl ethanolamide-d4 | LEA-d<sub>4</sub> | 18 | 2 | HMDB0012252 |
| Oleoyl ethanolamide-d4 | OEA-d<sub>4</sub> | 18 | 1 | HMDB0002088 |
| Palmitoyl ethanolamide-d5 | PEA-d<sub>5</sub> | 16 | 0 | HMDB0002100 |
| Stearoyl ethanolamide-d3 | SEA-d<sub>3</sub> | 18 | 0 | HMDB0013078 |

C – carbon chain length, C=C – number of carbon-carbon double bonds, HMDB – Human metabolome database.

### Instruments and apparatus

A centrifuge 5427 R from Eppendorf (Hamburg, Germany) was utilised for centrifugation during sample preparation. A SPD130DLX SpeedVac cold trap concentrator from Thermo Scientific (Midland, MI, USA) was also used for sample preparation.

The analytes were separated by liquid chromatography, followed by electrospray ionization (ESI) in positive mode and dMRM MS/MS detection. The analytical system comprised of Agilent 1290 Infinity UHPLC system coupled with an Agilent 6460 triple quadrupole (QqQ) mass spectrometer from Agilent Technologies Inc. (Santa Clara, CA, USA). The data were processed using Agilent MassHunter quantitative software (version 10.2).

### UHPLC-MS/MS method

The separation of the target analytes was carried out using Waters HSS T3 (1.8 μm, 2.1 × 100 mm) column, protected by C18 HSS T3 VanGuard pre-column (100Å, 1.8 µm, 2.1 mm × 5 mm) from Waters (Milford, Massachusetts, US.). The injection volume was 50 μL and the column temperature was set to 45°C. Mobile phase A was water with 0.1% formic acid, while mobile phase B was acetonitrile/isopropanol (v/v) with 0.1% formic acid. The flow rate was 0.4 mL min^-1^. The following gradient profile was employed, in positive mode analysis: From 0 to 1 minutes 60% A and 40% B, the percentage of A was then reduced to 20% between 1 to 2 minutes, for the next 7 minutes the percentage of A was reduced to 0% and then ramped up quickly to the starting conditions at 60% between 9 to 12 minutes. The total analysis time was 12 minutes. Instrument-dependent parameters for MS/MS were set as follows: the nitrogen drying gas flow rate was 12 L/min, with a temperature of 325 °C; the capillary voltage was 3500; the nebulizer pressure was maintained at 45 psi; the nitrogen sheath gas flow rate was 11.0 L/min, with a temperature of 325 °C. The dMRM was used for data collection, it monitored and selected the precursor product ion transitions of each analyte (Supplementary table 1). The data were processed using Agilent MassHunter quantitative software (version 10.2).

### Preparation of standards

An ISTD stock solution was prepared by dissolving each of the seven deuterated NAEs (Table 1) in pure organic solvent (MeOH, EtOH or ACN) to reach an adequate concentration of ∼1000 mg/L, and the samples were stored at –20°C. Seven calibration concentration levels were created from the ISTD stock solutions at 0.12, 0.48, 1.9, 7.8, 31.25, 125 and 500 ng/mL. A multistandard working solution that contains all the deuterated standards was prepared at a concentration of 10 mg/L for each standard in MeOH and stored at – 20°C.

### Sample extraction optimization

To evaluate extraction efficiency in human CSF samples in a broad range of polarities six different monophasic extraction solvents were compared: MeOH (extraction 1), ACN (extraction 2), ACN/Ace 1:1 (v/v) (extraction 3), CHCl_3_/ MeOH/H_2_O 2:5:2 (v/v/v) (extraction 4), IPA/ACE/ H_2_O 3:3:2 (v/v/v) (extraction 5), and EtOH (extraction 6).

50 μL of CSF was aliquoted in an Eppendorf tube. 10 μL of the ISTD mixture to each CSF sample using six different extraction solvents, individually to each CSF aliquot. The CSF samples were vortexed for 5 seconds, shaken at 1500 RPM for 5 minutes at 4 ° C and vortexed again for 5s before being centrifuged at 12,700 RPM for 5 min at 4 ° C. The supernatants (450 μL) were evaporated to dryness with a SpeedVac cold trap concentrator. All dried extracts were reconstituted in a 50 μL mixture of ethanol and toluene 9:1 (v/v). The reconstituted samples were analyzed by UHPLC-MS/MS. This procedure was carried out in six replicates and MS peak area were measured for the seven NAEs.

Recovery and precision were determined by measuring seven ISTDs. Recovery % was evaluated by comparing the CSF extracted ISTDs with the same ISTDs in buffer. The precision was defined as the relative standard deviation (RSD) in percent. The inter-day recovery and precision were calculated based on data collected from CSF and standards at three concentration levels high 90, medium 9 and low 0.9 ng mL^-1^, repeated over three days. QC samples at each concentration level were run in six replicates. Matrix effect was evaluated at three concentration levels high 90, medium 9 and low 0.9 ng mL^-1^. The matrix interference was evaluated based on mean peak area ratios of CSF samples post extraction to standards in buffer. LOD and LOQ were calculated according to the US Food and Drug Administration (FDA) recommendations (19). The LOD and LOQ were calculated as three and ten times, respectively, of the standard deviation divided by the estimated concentration based on the calibration curves (20). Linearity was investigated by examining the calibration curve of ISTDs at seven concentration levels, fitted to a linear regression model. Visualizations were carried out in R version 4.3.0.

The best extraction in terms of recovery and precision was using ACN/Ace and used for the n=33 human CSF samples. The method used for these samples is described in the UHPLC-MS/MS method section. These were run in triplicate and assumed to represent amounts in CSF in healthy individuals.

## Results and Discussion

To measure the CSF NAE concentrations in healthy individuals, we considered the following steps.

### Recovery of NAEs in CSF samples

The extraction efficiency of six different solvents was compared by assessing both extraction recovery and precision of the seven ISTDs (Fig.1). The extraction recovery in MeOH was within the range of 95-132%, 87-115% in ACN, 65-95% in CHCl_3_/MeOH/H_2_O, 81-106% in ACN/Ace, 52-144% in IPA/ACN/H_2_O, and 51-146% in EtOH. The acceptable recommended range for extraction efficiency is between 70-120% (21). This parameter was satisfied by three out of six solvent mixtures, ACN, CHCl_3_/MeOH/H_2_O and ACN/Ace.

**Fig. 1.**
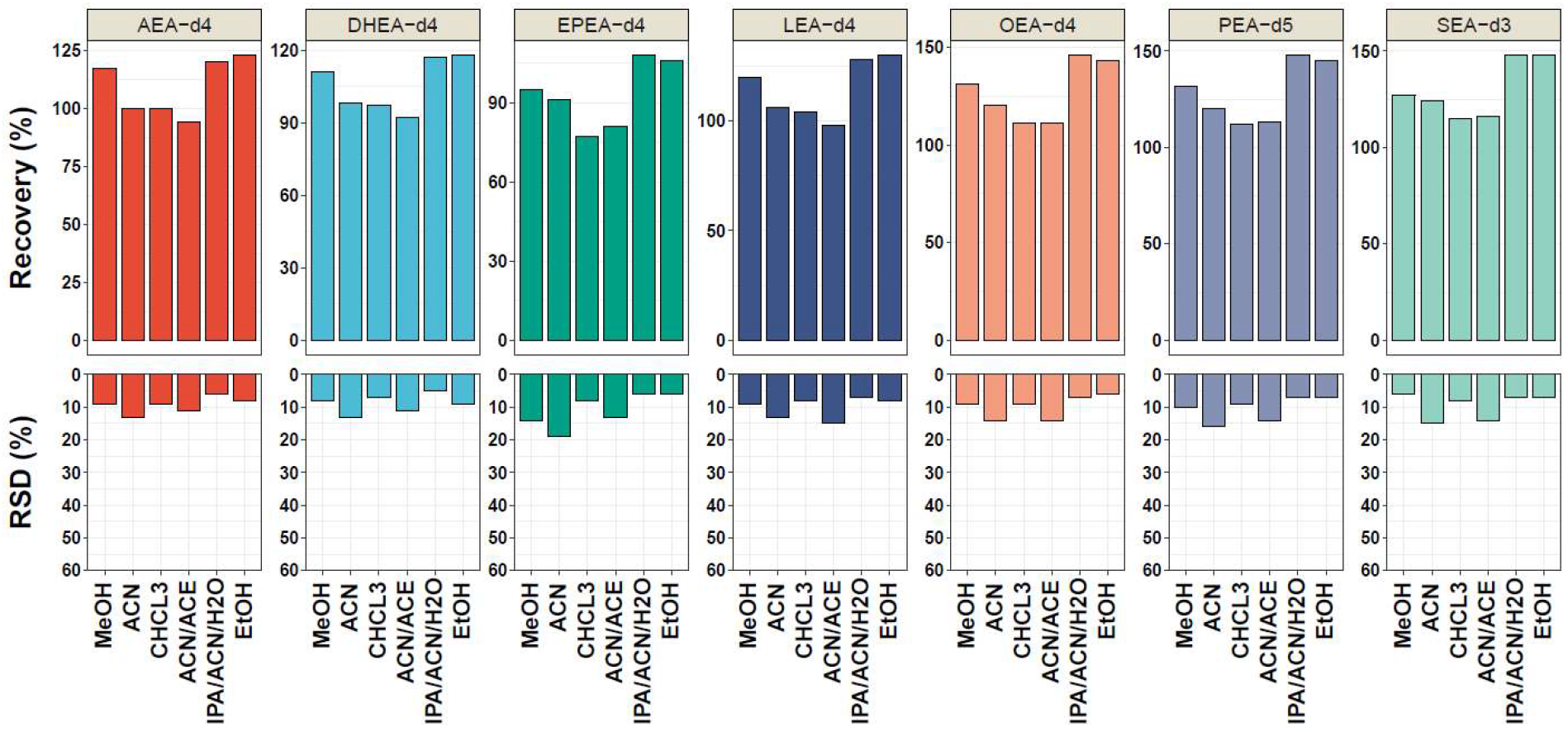
Extraction recovery and precision (%RSD) of the target NAEs (*n=7*) tested in six different monophasic solvents. MeOH = methanol. ACN = acetonitrile. CHCL3 = chloroform/methanol/water. ACN/Ace = acetonitrile/acetone. IPA/ACN/H2O = isopropanol/acetonitrile/water. EtOH = ethanol.

The precision expressed as RSD was in the range between 6-12% in MeOH, 14-21% in ACN, 41-47% in CHCl_3_/MeOH/H_2_O, 15-20% ACN/Ace, 7-9% in IPA/ACN/H_2_O, and 5-9% in EtOH. Four out of six solvents, MeOH, ACN/Ace, IPA/ACN/H_2_O and EtOH, satisfied these requirements.

The conditions for determining the most efficient extraction were met by ACN/Ace solvent mixture which was slightly more precise than ACN alone.

The recovery results for CHCl_3_/MeOH/H_2_O were similar to those observed by Reichl et al. (22) when utilizing a similar method for the liquid-liquid extraction of endogenous lipids in CSF. Similarly, ACN extraction recovery was within the accepted range. One study reported that acetone tends to co-precipitate lipids with proteins (23) and another that ACN might be best to precipitate most of the proteins in the sample (24). The combination of ACN/Ace solvent mixture showed acceptable recovery and precision (Fig.1) and since it was slightly better in precision than ACN alone it was chosen for the human CSF samples.

### Inter-day recovery and precision

The mean of the accuracy values reported as inter-day recoveries are listed in Table 2. All target NAEs showed inter-day recovery values ranging between 80-120% which was in the recommended range (19). The inter-day precision, reported as %RSD, varied between 3% to 12% which is within the recommended 20%. This suggests that the variability and repeatability of the results were within the tolerance limits.

**Table 2.** Mean inter-day recovery and precision (%RSD) of the target NAEs at three different concentration levels (low = 0.9, mid = 9, high = 90) ng mL^-1^ extracted with ACN/Ace over a period of three days.

| Analyte | Mean Inter-day Recovery (%) ± RSD (%) | | | | | | | | |
| --- | --- | --- | --- | --- | --- | --- | --- | --- | --- |
|  | Day 1 |  |  | Day 2 |  |  | Day 3 |  |  |
|  | Low | Medium | High | Low | Medium | High | Low | Medium | High |
| AEA-d<sub>4</sub> | 92 ± 9 | 102 ± 7 | 104 ± 7 | 97 ± 7 | 104 ± 6 | 97 ± 5 | 104 ± 15 | 103 ± 7 | 96 ± 4 |
| DHEA-d<sub>4</sub> | 90 ± 6 | 106 ± 8 | 107 ± 7 | 97 ± 10 | 104 ± 7 | 97 ± 4 | 101 ± 10 | 108 ± 7 | 98 ± 4 |
| EPEA-d<sub>4</sub> | 86 ± 7 | 100 ± 7 | 102 ± 8 | 90 ± 6 | 101 ± 7 | 93 ± 5 | 105 ± 11 | 104 ± 8 | 96 ± 4 |
| LEA-d<sub>4</sub> | 97 ± 6 | 109 ± 7 | 107 ± 7 | 106 ± 6 | 111 ± 6 | 103 ± 4 | 118 ± 14 | 112 ± 8 | 101 ± 4 |
| OEA-d<sub>4</sub> | 98 ± 6 | 109 ± 8 | 104 ± 9 | 117 ± 4 | 120 ± 7 | 109 ± 5 | 112 ± 10 | 114 ± 6 | 101 ± 4 |
| PEA-d<sub>5</sub> | 102 ± 8 | 113 ± 7 | 103 ± 10 | 102 ± 6 | 113 ± 7 | 103 ± 3 | 114 ± 12 | 118 ± 8 | 105 ± 4 |
| SEA-d<sub>3</sub> | 100 ± 5 | 114 ± 7 | 106 ± 10 | 108 ± 4 | 114 ± 5 | 105 ± 4 | 115 ± 9 | 117 ± 7 | 106 ± 5 |

Our results were in accordance to a study by Castillo-Petnado et al. (15) reporting similar inter-day-variation. These values were also achieved with smaller diameter columns as reported by Kantae et al. (13) using nano-flow LC-MS and with micro-flow LC-MS in a full quantification (25).

### Matrix effect

The matrix effect results as listed in supplementary table 2 and ranged between 85-100%. If the response in the matrix was suppressed or enhanced by more than 20% it was concluded that the results were affected by the matrix (20). No significant interference from the matrix was observed. The variability of the matrix interference results was within 20% RSD for all the NAEs. A study by Aydin et al. (26) reported slight ion enhancement using a CSF surrogate which was not the case in our case with human CSF samples.

### Limit of detection, limit of quantification and linearity

The LODs of the NAEs extracted from CSF and NAEs added buffer ranged between 0.101-0.154 ng mL^-1^ and 0.023-0.071 ng mL^-1^, respectively. The LOQs of the analytes extracted from CSF and in buffer ranged from 0.338 - 0.515 ng mL^-1^ and 0.077-0.237 ng mL^-1^, respectively. The calibration curve was fitted by a linear regression, the coefficient of determination (R^2^) was 0.99 (Table 3). The LODs and LOQs were deemed satisfactory for measuring the CSF samples.

**Table 3.** Limit of detection (LOD), limit of quantification (LOQ) and coefficient of determination (R^2^) values of target analytes (*n=7*) measured over a period of three days at 0.9 ng mL^-1^.

| Analyte | $R^2$ | Standard | | CSF | |
| --- | --- | --- | --- | --- | --- |
|  |  | LOD (ng/ml) | LOQ (ng/ml) | LOD (ng/ml) | LOQ (ng/ml) |
| AEA-d<sub>4</sub> | 0.998 | 0.064 | 0.212 | 0.152 | 0.505 |
| DHEA-d<sub>4</sub> | 0.996 | 0.066 | 0.222 | 0.123 | 0.409 |
| EPEA-d<sub>4</sub> | 0.999 | 0.064 | 0.214 | 0.115 | 0.384 |
| LEA-d<sub>4</sub> | 0.995 | 0.040 | 0.132 | 0.154 | 0.515 |
| OEA-d<sub>4</sub> | 0.999 | 0.035 | 0.116 | 0.112 | 0.374 |
| PEA-d<sub>5</sub> | 0.999 | 0.062 | 0.205 | 0.144 | 0.481 |
| SEA-d<sub>3</sub> | 0.996 | 0.023 | 0.077 | 0.101 | 0.338 |

The LOQ of this method for OEA-d_4_ (0.374 ng mL^-1^) which was quite sensitive. The main proposed advantage is that UHPLC is an alternative approach when analysing low abundance compounds in CSF with restricted volumes. Moreover, while it is clear that previous developed methods are robust and sensitive, it is worth noting that this method showed equivalent sensitivity to specialized nano HPLC.

### NAEs and PFAMs in human CSF from healthy participants

The developed method was applied to CSF samples from 33 healthy individuals, women (n=19) and men (n=14). A profile of 17 molecules endogenous to the human CSF were measured Their concentrations levels were within 0.1 – 1000 ng mL^-1^ (Fig.2).

**Fig. 2.**
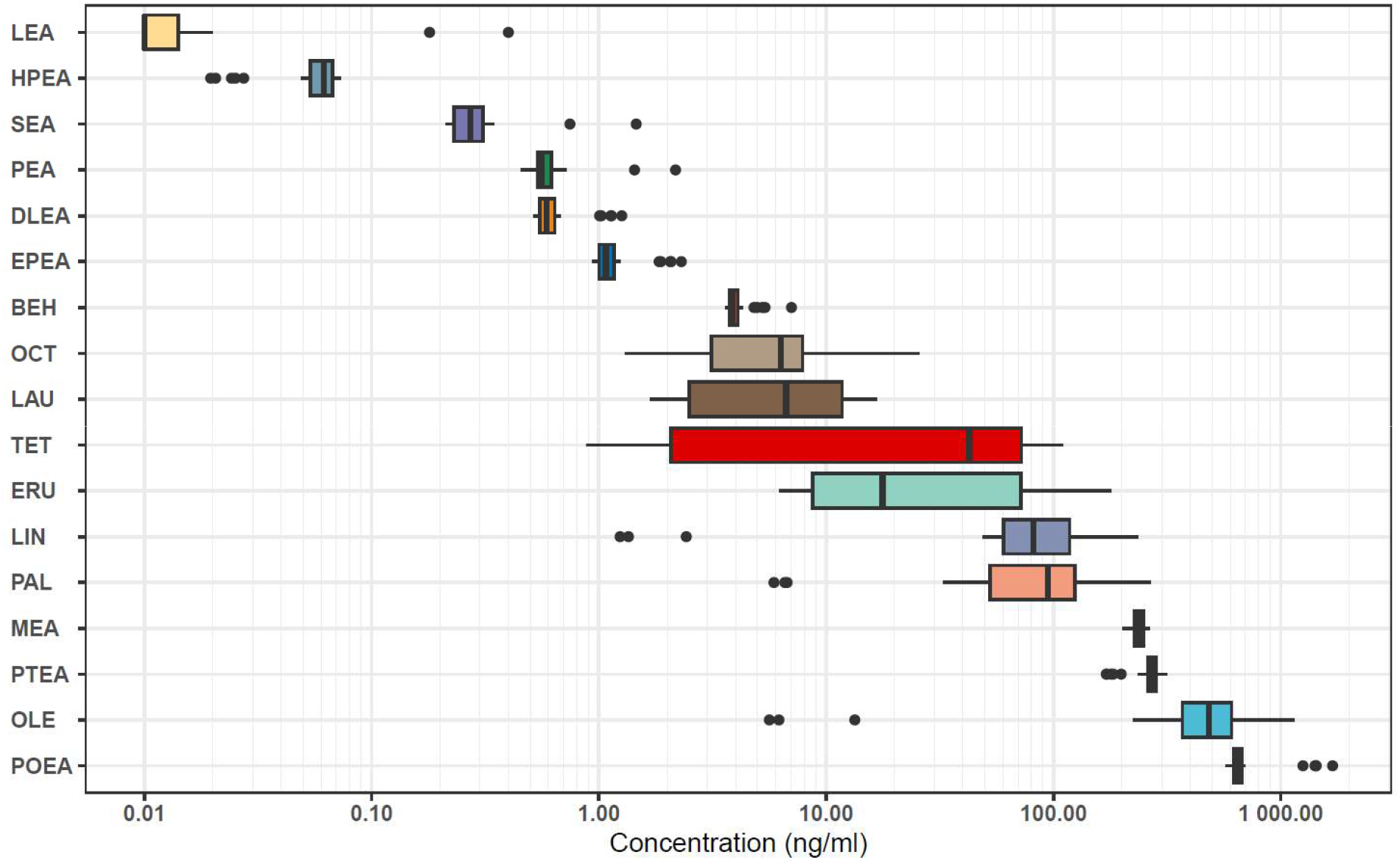
Concentration (ng mL^-1^) of 17 endogenous cannabinoid-like molecules in human CSF from 33 healthy individuals. (LEA - Linoleoyl ethanolamide; HPEA - Heptadecanoyl Ethanolamide; SEA - Stearoyl ethanolamide; PEA - Palmitoyl ethanolamide; DLEA - Dihomolinolenoyl ethanolamide; EPEA - Eicosatrienoic Acid Ethanolamide; BEH - Behenamide; OCT - Octadecanamide; LAU - Lauramide; TET - Tetradecanamide; ERU - Erucamide; LIN - Linoleamide; PAL - Palmitamide; MEA - Myristic monoethanolamide; PTEA - Pentadecanoyl Ethanolamide; OLE - Oleamide; POEA - Palmitoleoyl Ethanolamide).

In healthy participants we were able to identify seventeen small fatty amide molecules with potential neuroactive functions. An interesting observation that oleamide levels could be found in CSF at a very high range of concentrations up to 1000 ng mL^-1^. One study reports the concentration range of oleamide in plasma between 20 – 250 ng mL^-1^ (15). The difference in concertation between plasma and CSF could be due to oleamide synthesis which occurs in brain microglia (27). However, further research is required to confirm where oleamide synthesis occurs in humans.

A limitation of our method is that anandamide (AEA), also known as arachidonoyl ethanolamine was not detected in CSF with our method. This is an important molecule and its primary site synthesis is in the brain (28) however, some studies have reported intracellular stores of anandamide outside of the synaptic vesicles (29). Additionally, adipocytes have been identified as a novel site of anandamide synthesis (28). Anandamide is thought to induce anti-inflammatory responses (3,30) and play a role in sensory nerve fibers (8,31). The presence of extracellular anandamide transporters suggests that it could also be an endocrine messenger (32). It was detected in CSF using nano-LC technology in human CSF, to be on average 1.6 picomolar (13).

On the other hand, oleamide was detected and is also primarily localized within the brain microglia (27). Oleamide is detectable in human plasma and individuals diagnosed with Alzheimer’s disease (AD) can exhibit elevated levels within blood and exosomes (33,34). Its modulation could be related to sedative effects or sleep-inducing effects, maybe preventing hyperactivity in neurons (35).

## Conclusion

In this study, a UHPLC-MS/MS method for measuring NAEs and PFAMs in human CSF was shown. A solvent mixture of ACN/Ace yielded good extraction recovery results with good precision. The robustness and sensitivity of the method have been established by calculating the LOD and LOQ values, which were deemed satisfactory. Furthermore, the method can be applied to human CSF samples to better understand the biological purpose of neuroactive molecules.

## Abbreviations

(ACN): Acetonitrile
(CSF): cerebrospinal fluid
(dMRM): dynamic multiple reaction monitoring
(ESI): electrospray ionization
(FA): formic acid
(FDA): food and drug administration
(GC-MS): gas chromatography mass spectrometry
(IPA): isopropanol
(LOD): limit of detection
(LOQ): limit of quantification
(MeOH): methanol
(NAE): n-acylethanolamine
(PFAM): primary fatty amide
(RSD): relative standard deviation
(ISTD): internal standard
(QqQ): triple quadrupole
(UHPLC-MS): ultra-high performance liquid chromatography mass spectrometry
(AEA): Anandamide
(BEH): Behenamide
(DLEA): Dihomolinolenoyl ethanolamide
(DHEA): Docosahexaenoyl ethanolamide
(EPEA): Eicosapentaenoyl ethanolamide
(ERU): Erucamide
(HPEA): Heptadecanoyl Ethanolamide
(LAU): Lauramide
(LIN): Linoleamide
(LEA): Linoleoyl ethanolamide
(MEA): Myristic monoethanolamide
(OCT): Octadecanamide
(OLE): Oleamide
(OEA): Oleoyl ethanolamide
(PAL): Palmitamide
(POEA): Palmitoleoyl Ethanolamide
(PEA): Palmitoyl ethanolamide
(PTEA): Pentadecanoyl Ethanolamide
(SEA): Stearoyl ethanolamide
(TET): Tetradecanamide

## Declarations

### Ethics approval

The current study received approval from the Ethics Committee of the Danish Capital Region (Journal No.: H-21051757) and The Danish Data Protection Agency (Journal No.: P-2022-97). Informed consent was obtained from all subjects selected for the study prior to their participation at the Copenhagen Memory Clinic, Copenhagen University Hospital, Rigshospitalet. A total of 33 subjects participated in the study as non-demented healthy controls (NDHC).

### Availability of data and materials

The datasets generated and/or analysed during the current study are not publicly available but are available from the corresponding author on reasonable request.

## Acknowledgements

The authors thank participants in this study. We would also like to thank the technical staff across all the affiliations for their assistance in the laboratory.

## Funding

This work was supported by a grant from Lundbeck Foundation, Grant No.: R344-2020-989. Additional contributors were the Absalon Foundation of May 1^st^, 1978, and Toyota-Fonden Denmark to AHS and SGH.

## Competing Interests

The authors declare no competing interests.

## Authors contributions

CLQ and KH conceived and planned, KH and VI performed experiments and LC-MS analysis. KH, VI, AW analyzed the data and performed statistical analyses. AW, VI and CLQ wrote the manuscript. All authors provided critical feedback and helped to revise the manuscript.

## Consent to participate

Not applicable.

## Consent for publication

Not applicable.

## Notes

### Competing Interest Statement

The authors have declared no competing interest.

### Summary of Updates

This version of the manuscript has been revised to highlight the methods capabilities on N-acylethanolamines

## References

1. Hansen HS, Moesgaard B, Petersen G, & Hansen HH. Putative neuroprotective actions of Nacyl-ethanolamines. Pharmacol Ther. 2002;95(2):119–26. doi: 10.1016/S0163-7258(02)00251-6

2. Herrera MI, Kölliker-Frers R, Barreto G, Blanco E, & Capani F. Glial modulation by Nacylethanolamides in brain injury and neurodegeneration. Front Aging Neurosci. 2016;8(APR):1–10. doi: 10.3389/fnagi.2016.00081

3. Mock ED, Gagestein B, & van der Stelt M. Anandamide and other N-acylethanolamines: A class of signaling lipids with therapeutic opportunities. Prog Lipid Res [Internet]. 2023;89(July 2022):101194. doi: 10.1016/j.plipres.2022.101194

4. Hill MN, Miller GE, Carrier EJ, Gorzalka BB, & Hillard CJ. Circulating endocannabinoids and N-acyl ethanolamines are differentially regulated in major depression and following exposure to social stress. Psychoneuroendocrinology. 2009;34(8):1257–62. doi: 10.1016/j.psyneuen.2009.03.013

5. Ogawa S, Hattori K, Sasayama D, Yokota Y, Matsumura R, Matsuo J et al. Reduced cerebrospinal fluid ethanolamine concentration in major depressive disorder. Sci Rep. 2015;5:1–8. doi: 10.1038/srep07796

6. Esposito E, Cordaro M, & Cuzzocrea S. Roles of fatty acid ethanolamides (FAE) in traumatic and ischemic brain injury. Pharmacol Res [Internet]. 2014;86:26–31. doi: 10.1016/j.phrs.2014.05.009

7. Fernández-Irigoyen J, Cartas-Cejudo P, Iruarrizaga-Lejarreta M, & Santamaría E. Alteration in the cerebrospinal fluid lipidome in Parkinson’s disease: A post-mortem pilot study. Biomedicines. 2021;9(5). doi: 10.3390/biomedicines9050491

8. Di Marzo V, Fontana A, Cadas H, Schinelli S, Cimino G, Schwartz JC et al. Formation and inactivation of endogenous cannabinoid anandamide in central neurons. Nature. 1994;372(6507):686–91. doi: 10.1038/372686a0

9. Van Der Stelt M, Veldhuis WB, Van Haaften GW, Fezza F, Bisogno T, Bär PR et al. Exogenous anandamide protects rat brain against acute neuronal injury in vivo. J Neurosci. 2001;21(22):8765–71. doi: 10.1523/jneurosci.21-22-08765.2001

10. Clapper JR, Moreno-Sanz G, Russo R, Guijarro A, Vacondio F, Duranti A et al. Anandamide suppresses pain initiation through a peripheral endocannabinoid mechanism. Nat Neurosci. 2010;13(10):1265–70. doi: 10.1038/nn.2632

11. Brown I, Cascio MG, Wahle KWJ, Smoum R, Mechoulam R, Ross RA et al. Cannabinoid receptor-dependent and -independent anti-proliferative effects of omega-3 ethanolamides in androgen receptor-positive and -negative prostate cancer cell lines. Carcinogenesis. 2010;31(9):1584–91. doi: 10.1093/carcin/bgq151

12. Kim M, Snowden S, Suvitaival T, Ali A, Merkler DJ, Ahmad T et al. Primary fatty amides in plasma associated with brain amyloid burden, hippocampal volume, and memory in the European Medical Information Framework for Alzheimer’s Disease biomarker discovery cohort Lutz Fr €. Alzheimer’s Dement. 2019;15:817–27. doi: 10.1016/j.jalz.2019.03.004

13. Kantae V, Ogino S, Noga M, Harms AC, Van Dongen RM, Onderwater GLJ et al. Quantitative profiling of endocannabinoids & related N-Acylethanolamines in human CSF using nano LC-MS/MS. J Lipid Res [Internet]. 2017;58(3):615–24. doi: 10.1194/jlr.D070433

14. Sakka L, Coll G, & Chazal J. Anatomy and physiology of cerebrospinal fluid. Eur Ann Otorhinolaryngol Head Neck Dis [Internet]. 2011;128(6):309–16. doi: 10.1016/j.anorl.2011.03.002

15. Castillo-Peinado LS, López-Bascón MA, Mena-Bravo A, Luque de Castro MD, & Priego-Capote F. Determination of primary fatty acid amides in different biological fluids by LC– MS/MS in MRM mode with synthetic deuterated standards: Influence of biofluid matrix on sample preparation. Talanta [Internet]. 2019;193(September 2018):29–36. doi: 10.1016/j.talanta.2018.09.088

16. Hooshmand K, Xu J, Simonsen AH, Wretlind A, de Zawadzki A, Sulek K et al. Human Cerebrospinal Fluid Sample Preparation and Annotation for Integrated Lipidomics and Metabolomics Profiling Studies. Mol Neurobiol [Internet]. 2024;61(4):2021–32. doi: 10.1007/s12035-023-03666-4

17. Sultana T, & Johnson ME. Sample preparation and gas chromatography of primary fatty acid amides. J Chromatogr A. 2006;1101(1–2):278–85. doi: 10.1016/j.chroma.2005.10.027

18. Lee HJ, Kremer DM, Sajjakulnukit P, Zhang L, & Lyssiotis CA. A large-scale analysis of targeted metabolomics data from heterogeneous biological samples provides insights into metabolite dynamics. Metabolomics [Internet]. 2019;15(7):1–13. doi: 10.1007/s11306-019-1564-8

19. FDA. Bioanalytical Method Validation. U.S. Department of Health and Human Services Food and Drug Administration Center for Drug Evaluation and Research (CDER) Center for Veterinary Medicine (CVM). 2018. doi: 10.1201/9780203026427-15

20. Hooshmand K, & Fomsgaard IS. Analytical methods for quantification and identification of intact glucosinolates in arabidopsis roots using lc-qqq(Lit)-ms/ms. Metabolites. 2021;11(1):1– 18. doi: 10.3390/metabo11010047

21. Steiner D, Krska R, Malachová A, Taschl I, & Sulyok M. Evaluation of Matrix Effects and Extraction Efficiencies of LC-MS/MS Methods as the Essential Part for Proper Validation of Multiclass Contaminants in Complex Feed. J Agric Food Chem. 2020;68(12):3868–80. doi: 10.1021/acs.jafc.9b07706

22. Reichl B, Eichelberg N, Freytag M, Gojo J, Peyrl A, & Buchberger W. Evaluation and optimization of common lipid extraction methods in cerebrospinal fluid samples. J Chromatogr B Anal Technol Biomed Life Sci [Internet]. 2020;1153(April):122271. doi: 10.1016/j.jchromb.2020.122271

23. Awad D, & Brueck T. Optimization of protein isolation by proteomic qualification from Cutaneotrichosporon oleaginosus. Anal Bioanal Chem. 2020;412(2):449–62. doi: 10.1007/s00216-019-02254-7

24. Periasamy P, Rajandran S, Ziegman R, Casey M, Nakamura K, Kore H et al. A simple organic solvent precipitation method to improve detection of low molecular weight proteins. Proteomics. 2021;21(2100152):1–12. doi: 10.1002/pmic.202100152.This

25. He B, Di X, Guled F, Harder AVE, van den Maagdenberg AMJM, Terwindt GM et al. Quantification of endocannabinoids in human cerebrospinal fluid using a novel micro-flow liquid chromatography-mass spectrometry method. Anal Chim Acta [Internet]. 2022;1210(April):339888. doi: 10.1016/j.aca.2022.339888

26. Aydin E, Cebo M, Mielnik J, Richter H, Schüle R, Sievers-Engler A et al. UHPLC-ESI-MS/MS assay for quantification of endocannabinoids in cerebrospinal fluid using surrogate calibrant and surrogate matrix approaches. J Pharm Biomed Anal. 2023;222(October 2022). doi: 10.1016/j.jpba.2022.115090

27. Mendelson WB, & Basile AS. The hypnotic actions of the fatty acid amide, oleamide. Neuropsychopharmacology. 2001;25(5):S36–9. doi: 10.1016/S0893-133X(01)00341-4

28. Gonthier MP, Hoareau L, Festy F, Matias I, Volenti M, Bès-Houtmann S et al. Identification of endocannabinoids and related compounds in human fat cells. Obesity. 2007;15(4):837–45. doi: 10.1038/oby.2007.581

29. Aso E, & Ferrer I. Cannabinoids for treatment of alzheimer’s disease: Moving toward the clinic. Front Pharmacol. 2014;5 MAR(March):1–11. doi: 10.3389/fphar.2014.00037

30. Decara J, Rivera P, López-Gambero AJ, Serrano A, Pavón FJ, Baixeras E et al. Peroxisome Proliferator-Activated Receptors: Experimental Targeting for the Treatment of Inflammatory Bowel Diseases. Front Pharmacol. 2020;11(May):1–18. doi: 10.3389/fphar.2020.00730

31. Du Q, Liao Q, Chen C, Yang X, Xie R, & Xu J. The Role of Transient Receptor Potential Vanilloid 1 in Common Diseases of the Digestive Tract and the Cardiovascular and Respiratory System. Front Physiol. 2019;10(August). doi: 10.3389/fphys.2019.01064

32. Maccarrone M, Dainese E, & Oddi S. Intracellular trafficking of anandamide: new concepts for signaling. Trends Biochem Sci. 2010;35(11):601–8. doi: 10.1016/j.tibs.2010.05.008

33. Gómez-Pascual A, Naccache T, Xu J, Hooshmand K, Wretlind A, Gabrielli M et al. Paired plasma lipidomics and proteomics analysis in the conversion from mild cognitive impairment to Alzheimer’s disease. Comput Biol Med [Internet]. 2024;108588. doi: 10.1016/j.compbiomed.2024.108588

34. Kim M, Snowden S, Suvitaival T, Ali A, Merkler DJ, Ahmad T et al. Primary fatty amides in plasma associated with brain amyloid burden, hippocampal volume, and memory in the European Medical Information Framework for Alzheimer’s Disease biomarker discovery cohort. Alzheimer’s Dement. 2019;15(6):817–27.

35. Lerdkrai C, Asavapanumas N, Brawek B, Kovalchuk Y, Mojtahedi N, Del Moral MO et al. Intracellular Ca2+ stores control in vivo neuronal hyperactivity in a mouse model of Alzheimer’s disease. Proc Natl Acad Sci U S A. 2018;115(6):E1279–88. doi: 10.1073/pnas.1714409115

