## Supplementary tables for "A UHPLC–MS/MS method for determination of N-acylethanolamines in cerebrospinal fluid"

**Supplementary table 1** MRM transitions and target specific MS parameters of target analytes (n=8) used for the UHPLC-MS/MS analysis method.

| Analyte | $Q_1$ [m/z] | $Q_3$ [m/z] | Transition | Fragment (v) | CE (v) |
|---|---|---|---|---|---|
| AEA-$d_4$ | 352.3 | 66.2 | Quantifier | 92 | 13 |
| | | 91.1 | Qualifier | 92 | 53 |
| DHEA-$d_4$ | 376.3 | 66.2 | Quantifier | 128 | 17 |
| | | 91.1 | Qualifier | 128 | 57 |
| EPEA-$d_4$ | 350.3 | 66.2 | Quantifier | 106 | 5 |
| | | 91.1 | Qualifier | 106 | 57 |
| LEA-$d_4$ | 324.2 | 62.2 | Quantifier | 106 | 13 |
| | | 67.2 | Qualifier | 106 | 45 |
| OEA-$d_4$ | 330.3 | 66.2 | Quantifier | 106 | 13 |
| | | 55.2 | Qualifier | 106 | 49 |
| PEA-$d_5$ | 305.3 | 62 | Quantifier | 128 | 13 |
| | | 57.2 | Qualifier | 128 | 37 |
| SEA-$d_3$ | 331.3 | 62.2 | Quantifier | 128 | 13 |
| | | 57.2 | Qualifier | 128 | 41 |

$Q_1$ – first quadrupole, $Q_3$ – third quadrupole, CE – collision energy.

**Supplementary table 2** Mean matrix effect of each target NAEs at three different concentration levels (low = 0.9, mid = 9, high = 90) ng $mL^{-1}$ extracted with ACN/Ace.

| Analyte | Mean Matrix Effect (%) ± RSD (%) | | |
|---|---|---|---|
| | Low | Medium | High |
| AEA-$d_4$ | 85 ± 4 | 88 ± 5 | 89 ± 2 |
| DHEA-$d_4$ | 91 ± 4 | 92 ± 4 | 94 ± 3 |
| EPEA-$d_4$ | 85 ± 2 | 89 ± 4 | 91 ± 3 |
| LEA-$d_4$ | 87 ± 3 | 90 ± 4 | 94 ± 2 |
| OEA-$d_4$ | 85 ± 3 | 89 ± 3 | 92 ± 3 |
| PEA-$d_5$ | 85 ± 4 | 88 ± 4 | 92 ± 3 |
| SEA-$d_3$ | 87 ± 3 | 90 ± 3 | 94 ± 3 |
